# A multilayer perspective for inferring spatial and social functioning in animal movement networks

**DOI:** 10.1101/749085

**Authors:** Johann Mourier, Elodie J. I. Lédée, David M. P. Jacoby

## Abstract

1. Animal movement patterns are increasingly analysed as spatial networks. Currently, structures of complex movements are typically represented as a single-layer (or monoplex) network. However, aggregating individual movements, to generate population-level inferences, considerably reduces information on how individual or species variability influences spatial connectivity and thus identifying the mechanisms driving network structure remains difficult.
2. Here, we propose incorporating the recent conceptual advances in multilayer network analyses with the existing movement network approach to improve our understanding of the complex interaction between spatial and/or social drivers of animal movement patterns.
3. Specifically, we explore the application and interpretation of this framework using an empirical example of shark movement data gathered using passive remote sensors in a coral reef ecosystem. We first show how aggregating individual movement networks can lead to the loss of information, potentially misleading our interpretation of movement patterns. We then apply multilayer network analyses linking individual movement networks (i.e. layers) to the probabilities of social contact between individuals (i.e. interlayer edges) in order to explore the functional significance of different locations to an animal’s ecology.
4. This approach provides a novel and holistic framework incorporating individual variability in behaviour and inter-individual interactions. We discuss how this approach can be used in applied ecology and conservation to better assess the ecological significance of variable space use by mobile animals within a population. Further, we argue that the uptake of multilayer networks will significantly broaden our understanding of long-term ecological and evolutionary processes, particularly in the context of information or disease transfer between individuals.

## INTRODUCTION

Animal movements are complex and dynamic, involving behavioural and environmental drivers that can interact both in space and time. Consequently, ecological landscapes can be connected in ways that contain different layers of complexity, reflecting how species utilise the landscape in different ways and potentially influencing the services they provide (Bélisle, 2005). There has been great interest in understanding how animals use their habitat and conversely how habitat drives movement patterns (Hussey et al., 2015; Kays, Crofoot, Jetz, & Wikelski, 2015). Furthermore, while animal mobility influences population structure and ecological connectivity, this process can be highly dynamic due to individual behavioural variability in addition to the social ecology of the species in question. To ensure that behavioural data can be used for applied conservation, a quantitative understanding that fully captures these complex processes is essential (Nathan et al., 2008).

Network theory provides a powerful set of tools to visualise, quantify and interpret complex patterns of animal movements, conceptualised as spatial networks. Spatial networks have been employed increasingly in animal movement ecology to investigate different ecological processes such as pathogen transmission, gene flow, information transfer or nutrient transfer between habitats (e.g. Robertson et al., 2018; Williams, Papastamatiou, Caselle, Bradley, & Jacoby, 2018) as they offer a means to quantify links between locations and habitats established by animal movements using distinct metrics. Ecologists are beginning to combine recent advances in tracking technologies with developments in spatial networks (see Jacoby & Freeman, 2016 for a review). To date, spatial networks have been derived from a variety of tracking technologies including radio tracking or satellite transmitters (e.g. Shimazaki et al., 2004) and most commonly acoustic telemetry (e.g. Jacoby, Brooks, Croft, & Sims, 2012; Lédée, Heupel, Tobin, Mapleston, & Simpfendorfer, 2016).

Like most networks, movement networks evolve over time and are often derived from a subsample of individuals from a population for which tracking data were collected. Most spatial and movement ecology studies that employ network analyses have been restricted to monolayer networks, in other words, to a single, aggregated network layer. In fact, the traditional approach has been to build individual movement networks, but then to either compare network metrics between individuals using traditional statistics or null models or to aggregate each individual network into a single graph representing a subgroup of animals (e.g. sex or species). Mapping out these complex systems as monolayer networks can lead to the loss of important information, not least the role of individual variability on patterns of population connectivity and mobility (Spiegel, Leu, Bull, & Sih, 2017). In addition, while aggregating or comparing individual movement networks can provide useful information on the relative importance of some nodes (i.e. locations) based on their centrality, these networks may be misleading for interpreting processes that are facilitated by sociality. For example, monolayer movement networks do not consider contact rates between individuals, that is they typically cannot differentiate between animals using the same location at the same or at different times and this can have important implications for the probabilities of disease or information transfer between individuals (Silk et al., 2019). Therefore, there remains a number of analytical challenges associated with how best to amalgamate data to draw population-level inferences or to divide movement network data to explore individual- and group-level variation in behaviours.

Multilayer networks are a class of networks introduced to model systems composed of multiple layers giving rise to the idea of a ‘networks of networks’, allowing us to differentiate between intralayer (horizontal) and interlayer (vertical) connectivity (Kivelä et al., 2014). The theoretical basis for multilayer networks has been refined across many different research fields, already leading to important insight, for example from transportation networks (Cardillo et al., 2013) to disease epidemics (Pilosof, Greenbaum, Krasnov, & Zelnik, 2017). As research on complex systems in ecology has developed, it has become increasingly important to move beyond simple graphs and investigate more complex, but more realistic frameworks, that account for the many potential sources of variability within the system. In multiplex networks, a particular form of multilayer network, the nodes within each network ‘layer’ are occurrences of the same entity; that is, all nodes are replicated on all layers where they are connected differently, either by different individuals, different species or by different modes of movement (e.g. walking, flying). The theory and mathematical developments for multilayer network analyses are now sufficient for upgrading traditional representations of networks in ecology, providing novel insights into the structure, functioning and dynamics of movement networks (Boccaletti et al., 2014; Kivelä et al., 2014).

Here, we present a new conceptual framework for incorporating individual movement trajectories with inter-individual contact patterns within a multilayer network object. Our approach considers layers made up of individual movement networks. These individuals might connect nodes - representing fixed monitoring locations that are present within each layer - differently in the different layers. All layers are then connected together by the probability of co-occurring (i.e. contact) at a given node. This approach provides a flexible framework that not only considers the spatial drivers of movement (intralayer edges) but also the temporal/social drivers of movement (interlayer edges). We then map intra and interlayer network metrics within a bivariate space to explore the proportional influence of spatial and social influence on the functional significance of a monitoring location. In order to illustrate the value of this approach, we analysed the movements of individual of two shark species to (1) demonstrate that aggregating individual layers into a single aggregated network can lead to the loss of important information and heterogeneity in the data and (2) to quantify the spatio-social significance of acoustic receiver locations to the ecology of marine predators occupying a coral reef ecosystem. Finally, we discuss the potential ecological applications of this framework in evolutionary ecology and conservation biology, in addition to exploring the possible future developments within this field.

## MATERIALS AND METHODS

### A multilayer representation of animal movements

A multilayer network consists of a set of nodes representing entities (e.g. locations in the case of spatial movement networks) and a set of layers representing the same spatial configuration of these locations (Fig. 1a). According to the definition of (Kivelä et al., 2014), a multilayer network *M* has a set of nodes *V* included inside layers. The multilayer network is constructed by assembling all combinations of layers L using a Cartesian product *L*_*1*_ x … x *L*_*d*_, with *d* = 0 for a monolayer network, *d* = 1 for one type of layering and *d* = 2 for two types of layering. In some layers, nodes might not always be connected. To indicate whether a node is present in a layer, a subset of *V* x *L*_*1*_ x … x *L*_*d*_ of all combinations is defined as *V*_*M*_ ⊆ *V* x *L*_*1*_ x … x *L*_*d*_ which contains only the node-layer combinations in which a node is present in the corresponding layer. The edge set *E*_*M*_ ⊆ *V*_*M*_ x *V*_*M*_ including both intralayer and interlayer edges encodes the connections between pairs of state nodes (i.e. nodes that exists on a specific layer). A multilayer network is then defined as a quadruplet *M* = (*V*_*M*_, *E*_*M*_, *V*, L). A multilayer network can be represented by an adjacency matrix, also called a ‘supra-adjacency matrix’, with intralayer edges on the diagonal blocks and interlayer edges on the off-diagonal blocks (Fig. 1b).

**Figure 1:**
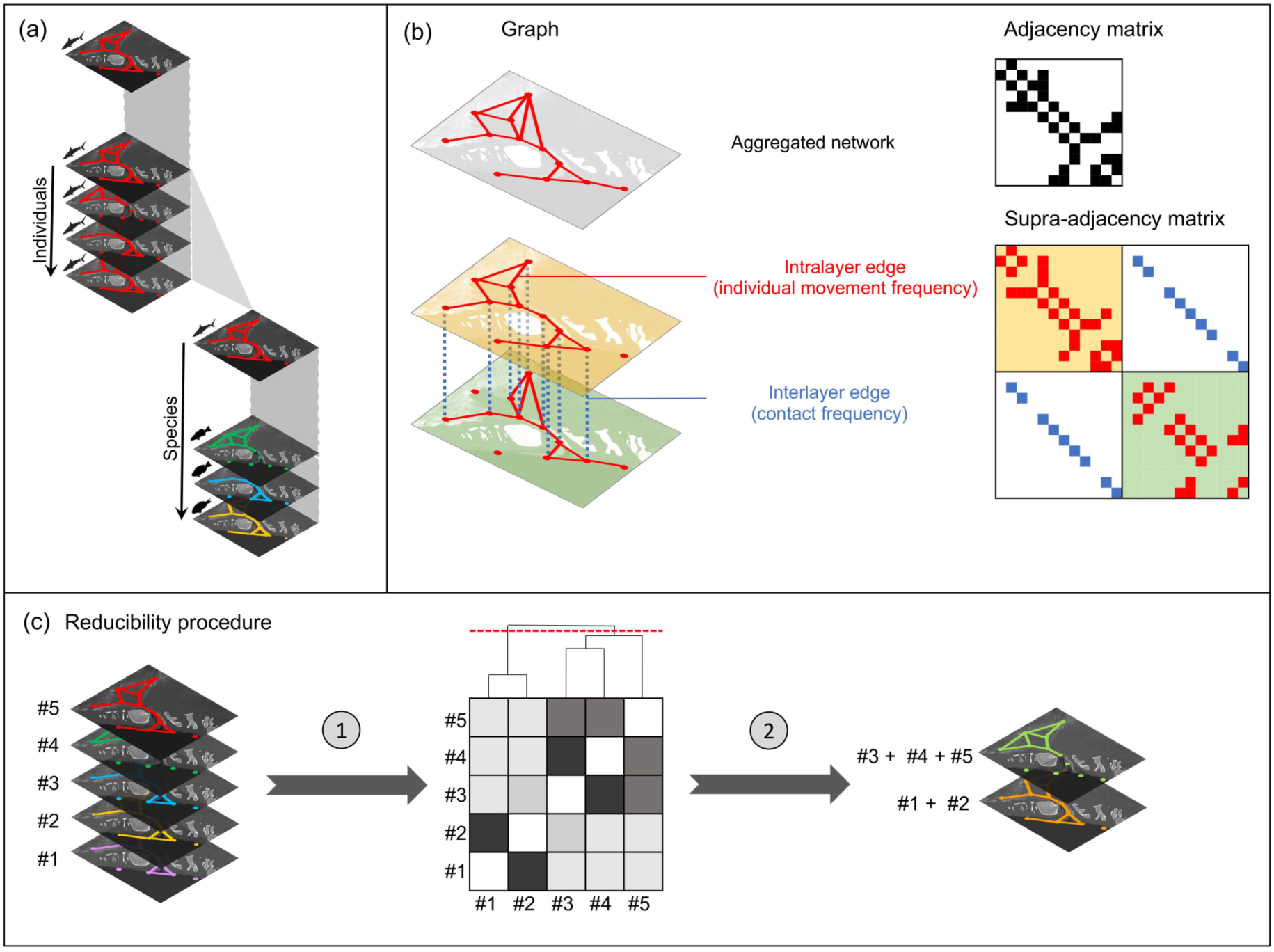
(a) The multi-layer nature of movement networks constituted of multiple individual movement networks from a same or different species. (b) The movement network of a species is often represented as an aggregated spatial network of many individuals from the same species, forming a monolayer network (grey). However, it can be represented as a multilayer network with each layer representing the movement network of an individual; here a multilayer network constituted of two individual movement networks (yellow and green layers) linked by interlayer edges which can represent frequency of contact between individuals at a particular node. The aggregated network is mathematically represented by an adjacency matrix and the multilayer network is represented by a supra-adjacency matrix including both intra and interlayer edges. (c) In order to optimize the amount of information in the multilayer movement network, a reducibility procedure can help removing redundant information about movement and merge layers of similar movement properties.

Here, the intralayer edges are built from the movements, and for a weighted network, the frequency of movements of each monitored individual. The interlayer edges link each node of all layers by a probability of contact or co-occurrence at a specific node (i.e. location). In fact, by linking nodes between layers by frequency of contact, we better delineate the temporal overlap that can improve our understanding of transfer of information, parasites or disease between layers (i.e. individuals) beyond simply considering spatial overlap. Incorporating both movement and contact in a multilayer framework will help identify the locations that are critical for such transfer processes.

Domenico, Nicosia, Arenas, & Latora (2015) describe how to aggregate multilayer networks in a way that attempts to minimize information loss and interlayer redundancy by maximising distinguishability between respective layers. This method of *reducibility* can provide a way to classify and aggregate individuals according to their similarity in movement network topology and keep only layers of dissimilar information for the overall multilayer network. Therefore, we may begin with all individuals as a separate layer but during pre-processing some of these individuals may be aggregated (Fig. 1c). Briefly, the reducibility procedure first computes the distance (based on quantum Jensen-Shannon divergence (Majtey, Lamberti, & Prato, 2005)) between all pairs of layers. Then hierarchical clustering of layers using the distance matrix is performed based on changes in relative entropy q(•). Finally, the best partition is determined as the one with the highest value of q(•). More details of this procedure can be found in (Domenico et al., 2015) with reducibility procedures carried out using the MuxViz toolkit (De Domenico, Porter, & Arenas, 2015).

### Node centrality and functionality

Network centrality metrics attempt to measure the importance of a node, an edge or a subgraph, and several monolayer centrality measures have been generalized to multilayer networks. Node centrality can be an important parameter in movement networks for identifying potential locations for targeted conservation efforts (Jacoby & Freeman, 2016). Node degree is a common centrality measure and is defined as the number of neighbouring nodes a node is directly connected to. In a multilayer network, degree heterogeneity might be present in two aspects: across the nodes in a layer and across layers in a node. The common approach infers the spatial centrality of a node either as a global trend resulting from the aggregation over all layers or as the correlated centrality tendencies of that node across all layers. However, both approaches do not consider interlayer edges, linking nodes between layers, and the centrality of a node which results from the spatial utilisation and preferences of all individuals might underestimate the node importance in connecting individuals through ecological processes, for example areas of importance for transmission of information or disease.

In our revised framework, we propose that the centrality of a node in a multilayer movement network would depend on two components: centrality based on the movement paths entering and leaving the node (i.e. from intralayer edges) which we refer to as “spatial centrality” and centrality based on the number and frequency of contacts between individuals occurring at a particular node (i.e. from interlayer edges) which we refer to as “social centrality”. Therefore, by considering the spatial and social components of movement within a population, functional centrality should be high when it has both high spatial centrality and high social centrality. To tease apart the relative importance of spatial and social centrality, we use the TOPSIS method (Technique for Order Preference by Similarity to an Ideal Solution; Huang, Keisler, & Linkov, 2011). This method is used to solve multiple criteria decisions by evaluating the alternatives by simultaneously measuring their distances to the Positive Ideal Solution (PIS) and to the Negative Ideal Solution (NIS). PIS is the most preferred solution (in our case the highest centrality) and NIS is the least preferred solution (the lowest centrality). The preference order from the TOPSIS method is then built according to the relative closeness of the alternatives to PIS, which is a scalar criterion that combines these two distance measures.

### Empirical example from acoustic tracking of sharks

Having outlined the rationale behind the integration of multilayer networks and multi-centrality approaches to movement ecology, we apply these methods to an empirical example of sharks tracked inside an acoustic array of monitoring stations. Movement data from grey reef sharks (*Carcharhinus amblyrhynchos*, Bleeker 1856) and blacktip reef sharks (*Carcharhinus melanopterus*, Quoy & Gaimard, 1824) at three offshore reefs (Heron, Sykes and One Tree) located in the southern Great Barrier Reef in Australia (Heupel, Lédée, & Simpfendorfer, 2018) were analysed using this methodology. The data were gathered as part of a long-term field study into the ecology of large predators along the coast of Australia. Open acoustic data were sourced from the Integrated Marine Observing System’s Animal Tracking Facility (IMOS ATF, www.imos.org.au). Specifically, we analyse the movement patterns of 24 individual grey reef sharks and 20 blacktip reef sharks fitted with long-life V16 acoustic transmitters. Grey reef sharks were monitored over a period of 13 months between 13^th^ April 2012 and 14^th^ May 2013 (34 047 detections) within an array of 48 VR2W hydrophone receivers (VEMCO, Halifax, Nova Scotia). Blacktip reef sharks were monitored over a period of 10 months between 31^st^ August 2013 and 16^th^ June 2014 (3 888 detections) within an array of 50 VR2W hydrophone receivers. Receivers detect acoustic transmitters at an approximate range of 270 m; each time transmitters are detected, the identification number, date and time are recorded by the receiver (more details can be found in Heupel et al. (2018)).

Applying network theory to acoustic telemetry data allowed the movement of sharks to be viewed as a system of connections, in which acoustic receivers are linked by shark movements (for further details on network theory and how it is applied to telemetry data, see Jacoby et al. (2012) and Mourier, Lédée, Guttridge, & Jacoby (2018)). To obtain individual movement networks for each tagged shark (i.e. layer), a square matrix was constructed consisting of counts of individual movements from one node (i.e. receiver) to another. Then the interlayer edges were determined by inferring the spatio-temporal co-occurrences at each node using a Variational Bayesian mixture model, GMMEvents of the detection time-series (Jacoby, Papastamatiou, & Freeman, 2016), so that a link was created between two layers at a node if the two individuals co-occurred at least once at this location.

In order to investigate the differences in functional network topology between the aggregated movement network and individual layers, we applied a reducibility procedure to the multilayer network (Domenico et al., 2015) using the platform MuxViz (De Domenico, Porter, et al., 2015). This procedure divides the aggregated network into clusters of different topologies (Figure 1c), in turn testing if important sub-structure is lost during the aggregation process. To explore differences in node centrality we first calculated node strength for the aggregated network (i.e. the sum of movements across all layers arriving to or leaving the focal node) for each node of all individual movement networks. We constructed an aggregated movement network by summing movements from all individuals between each pair of nodes and then calculated strength for each node of the aggregated network. As we were interested in comparing the relative spatial centrality of nodes, for all layers as well as for the aggregated network, we ranked nodes based on their strength providing a rank score ranging from 0 (lowest spatial centrality) to 1 (highest spatial centrality). Correlation between each pair of layers was inferred using Spearman’s correlation to assess the degree of similarity/dissimilarity between layers. We then determined the overall spatial centrality score by summing for each node its centrality rank in each layer and then ranking this score from 0 and 1 (i.e. the mean rank of each node across layers). Similarly, strength was also calculated for the social interactions by summing the number of co-occurrences at each node and then ranking them from 0 and 1. For each species, a bivariate analysis was conducted by plotting both centrality scores across the spatial and social axes with a view to exploring the functionality of nodes within the movement network. We then used the TOPSIS method to organise the nodes according to their distance to positive and negative ideal solutions. In our case, centrality was considered as a function of spatial and social centralities. We define the positive ideal solution corresponding to the condition where spatial and social centralities are maximised (note that in our case a 1:1 weight ratio of spatial and social components was used to rank nodes according to an equal consideration of their spatial and social score but other ratios could be used to give more weight to one component). TOPSIS scores are then ranked from 0 to 1 and correspond to the multilayer ranks integrating socio-spatial centralities. All analyses were conducted in R (R Core Team 2019).

## RESULTS

### Multilayer movement network of sharks in the Great Barrier Reef

We ranked the nodes of each individual layer of the weighted multilayer networks of grey reef and blacktip reef shark movements according to their weighted degree. We found a high level of variability between layers (Figs 2-3) with groups of positively correlated layers (i.e. similar individual spatial centralities). Node spatial centrality ranks of the aggregated network were significantly positively correlated with 20 layers (83% of individual layers) in grey reef sharks and just 10 layers (50% of individual layers) in blacktip reef sharks (Figs 2-3). Although the correlations are relatively high, node spatial centrality of the aggregated network seems more dissimilar to the spatial and social centralities of the multilayer and to the functional multilayer centralities (i.e. inferred from the TOPSIS method) (Figs 2-3). The reducibility procedure reduced structural redundancy in the movement networks from 24 layers to three layers in the grey reef shark network and from 20 layers to four layers in the blacktip reef shark network (Fig. 4). These clusters were deemed to contain topologically similar network properties but clearly illustrate the deferential space use across small clusters of individual sharks (Fig. 4).

**Figure 2:**
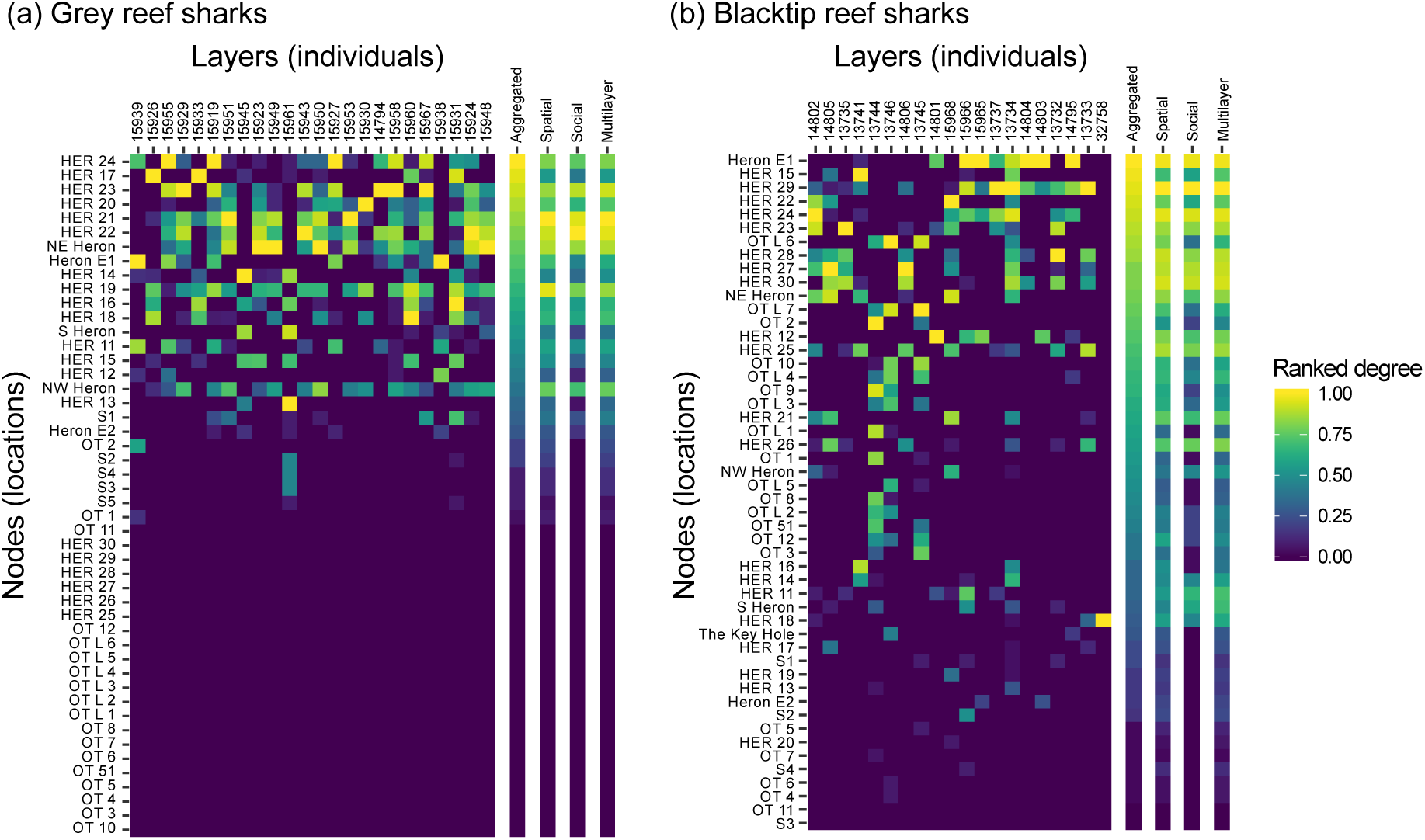
Heatmaps representing the ranks in centrality for each node in each layer for (a) grey reef sharks and (b) blacktip reef sharks. Note that the heatmaps are ordered by node centrality of the aggregated network in order to easily compare with the centrality of the different layers as well as the different methods (i.e. spatial, social and multilayer centralities). Ranks correspond to the weighted degree scores of each node ranging from 0 (blue; less central) to 1 (yellow; most central).

**Figure 3:**
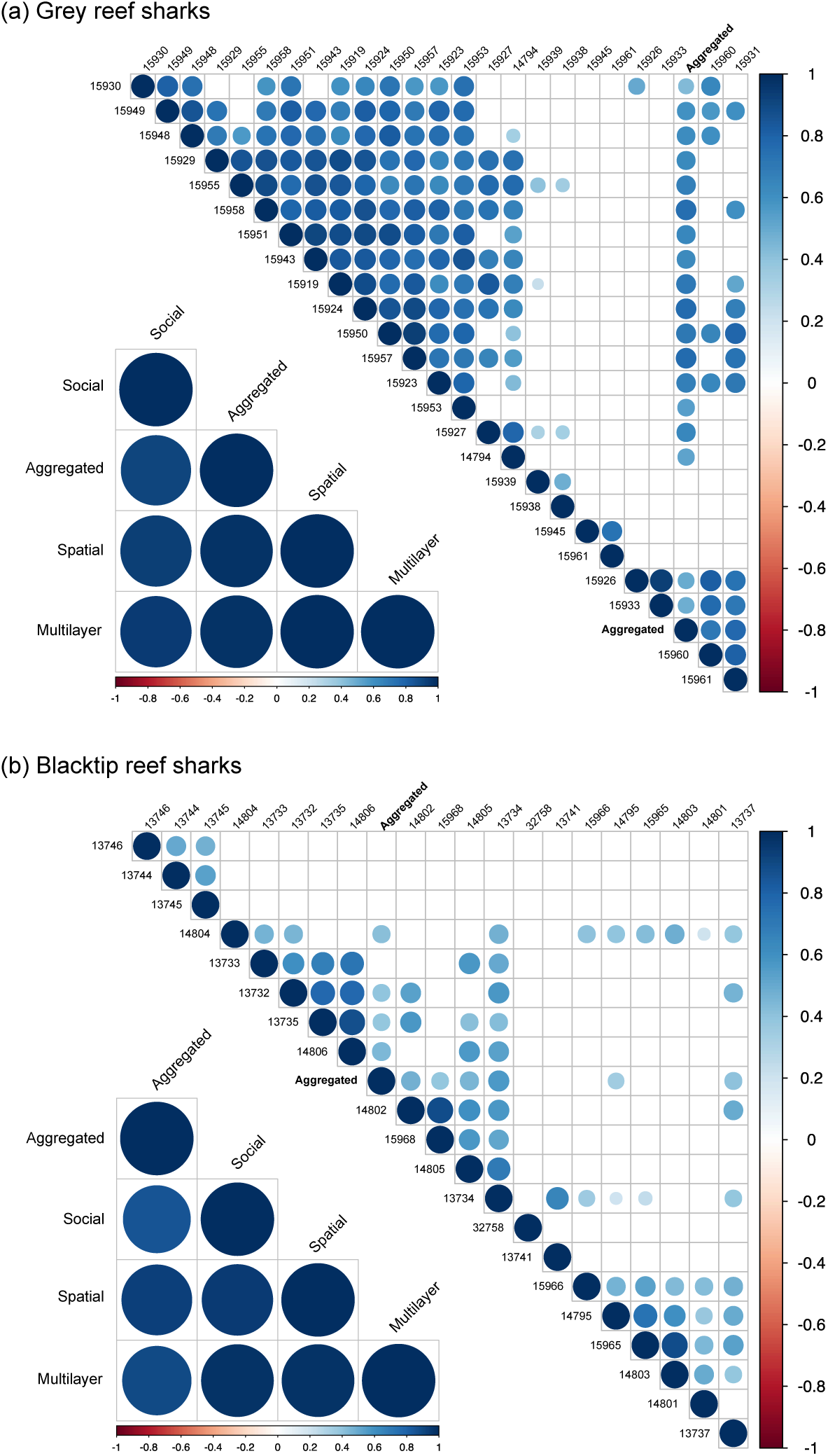
Correlation matrices of individual layers and the aggregated network (upper panel) and correlation matrices of the different global decision networks (including the aggregated, the average spatial “intralayer” and social “interlayer” networks as well as the multilayer network inferred from the TOPSIS analysis) for (a) grey reef sharks and (b) blacktip reef sharks. Colour intensity and the size of the circle are proportional to the correlation coefficients (positive correlation in blue and negative ones in red). Only significant correlations at the significant level of 0.01 are displayed, where blanks correspond to non-significant correlations.

**Figure 4:**
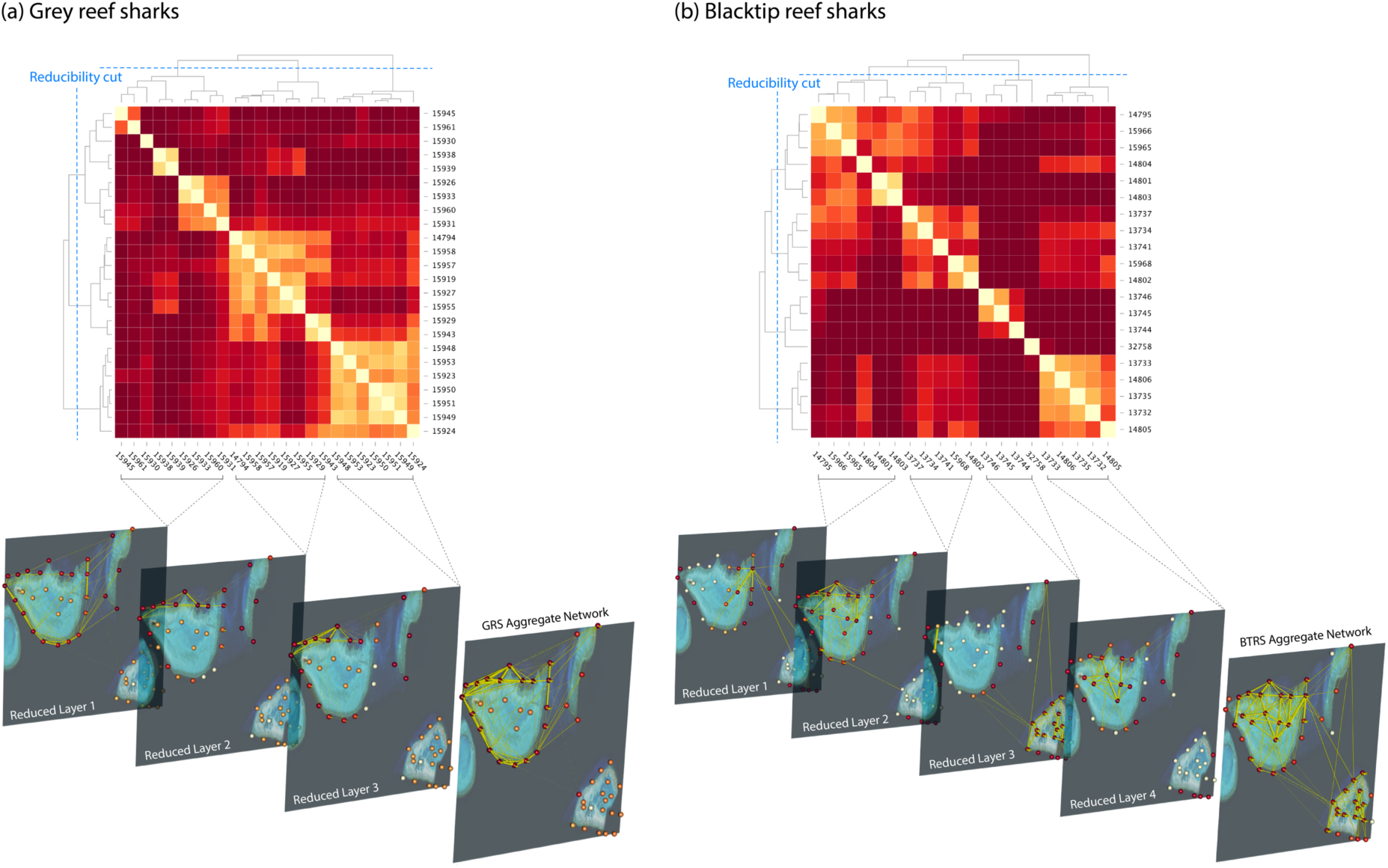
Reducibility procedure defining clusters of topologically similar individual movement networks for (a) grey reef sharks and (b) blacktip reef sharks. Movement networks within clusters were then combined as a layer within the multilayer network before being aggregated across layers. Nodes are coloured by degree centrality (yellow to red = low to high).

Incorporating social information into this picture, the bivariate analysis tends to show that spatial and social components of grey reef shark movement patterns are highly linked (Fig. 5a) with nodes of high spatial centrality being also of high social centrality. Conversely, spatial and social components of blacktip reef sharks are less associated in blacktip reef sharks (Fig. 5b) with nodes of moderate spatial centrality being of higher social centrality and nodes being more spatially than socially central. This analysis reveals the difference in functional importance of particular locations between species that use these areas in different ways ecologically.

**Figure 5:**
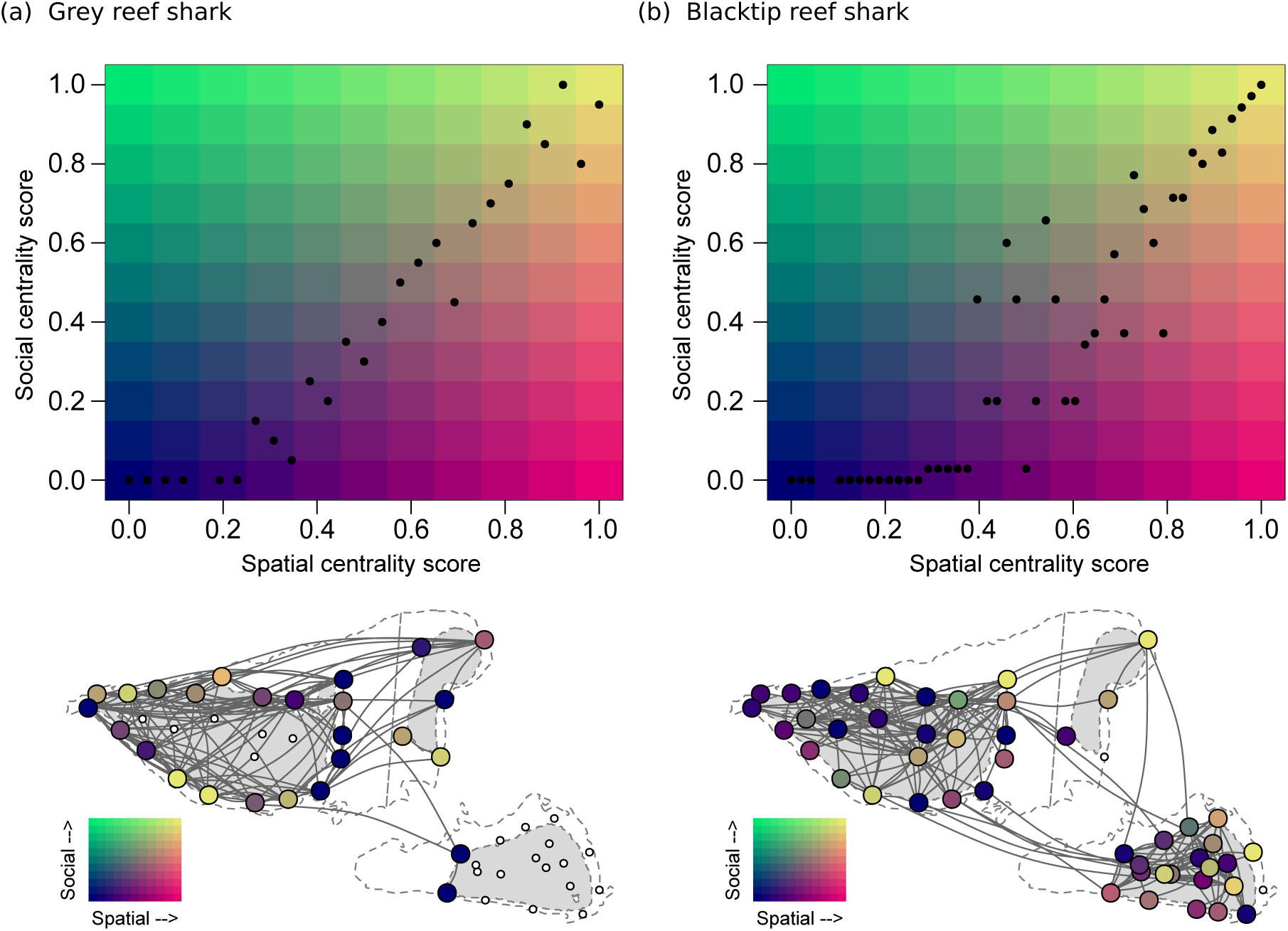
Bivariate map of nodes centrality ranks for (a) grey reef sharks and (b) blacktip reef sharks. Each black dot corresponds to a node located on a two-dimension map according to its social and spatial centrality score. Colours are representative of the functional centrality with nodes only important in terms of space use in pink, nodes in green only important in terms of sociality and nodes in yellow representative of both combined spatial and social centrality. Below is represented the corresponding aggregated movement network of both species where node colour is representative to their functional centrality score (i.e. their colour in the bivariate map) and edges represent the movements between nodes (for clearer visibility edge size is not proportional to edge weight).

## DISCUSSION

The application of multilayer methods are already gaining traction for exploring the ecological and evolutionary processes influencing animal populations (Silk, Finn, Porter, & Pinter-Wollman, 2018). In particular, studying interconnected networks could facilitate consideration of animal social systems in a broader ecological context, for example in understanding how habitat and movement networks influence the social structure of a population. Until now, multilayer approaches have been proposed independently in spatial and social network contexts (but see Manlove et al., 2018). Our method incorporates the recent conceptual advances in multilayer network analyses with the existing movement network approach to improve our understanding of the complex interaction between spatial and/or social drivers of animal movement patterns. It provides a novel and holistic framework linking social to spatial networks within a single network object for determining the functional significance of locations within an animal’s home range. These analyses demonstrate the importance of considering individual variability in movement networks to avoid the loss of important information on movement properties within the studied population. It also contributes to previous research demonstrating the importance of considering individual variability of movement and space-use for understanding the drivers of socio-spatial, ecological and evolutionary dynamics of animal populations (Spiegel et al., 2017). Furthermore, we explore this using a reducibility procedure to simplify and condense individuals that correlate strongly in their movement topologies, while reducing the amount of redundant information in the network (Domenico et al., 2015).

Our bivariate comparative analysis of the multilayer networks of two shark species demonstrates clear disparities in network topology that highlight ecological differences between the two species that are masked using single-layer network analyses (Heupel et al., 2018). The grey reef sharks showed a unimodal site functionality with spatial and social centralities correlating strongly with one another. Indeed, the most central nodes correspond to locations that are repeatedly used by multiple individuals. This is not surprising as this species is recognized as a central place forager (Mourier et al., 2016; Papastamatiou et al., 2018) forming large aggregations at specific locations where they are relatively resident. In contrast, the blacktip reef sharks showed a trimodal site functionality with some central nodes being simultaneously used for social and spatial purposes, while other nodes being independently used for social or spatial functions. Again, this is intuitive as they have been shown to develop complex socio-spatial behaviour (Mourier, Vercelloni, & Planes, 2012). Our framework confirms this property by showing that some locations are rarely visited (low spatial centrality) but specifically assigned to social activities (high social centrality) as many individuals co-occur when visiting these locations, while other locations are commonly visited but not for social purposes as individual rarely co-occur (i.e. no temporal overlap). These two examples illustrate the potential of this framework to improve our understanding of key ecological processes in animal movement ecology and the functional role of space utilisation and animal decision making.

### Potential applications of this framework

Applications of multilayer networks in spatial ecology and conservation are still scarce and some areas of general multilayer theory remain challenging or neglected. We anticipate that multilayer spatial networks might prove extremely valuable in conservation. Network theory has been increasingly applied in conservation in recent years, especially for identifying the centrality of certain locations in animal connectivity (Galpern, Manseau, & Fall, 2011). Depending on the species in question and the aims of the study, the relative importance of spatial and social components might differ. For example, in conservation, one might aim to identify locations at which most movements converge to manage areas of high flow, and therefore the spatial component might be of more importance. However, in certain cases, managers might aim to target areas where animals predominantly socialise (e.g. for mating or refuging purposes), and in this regard the social component would matter more. Finally, conservationists might aim to identify areas that are important for parasite/disease transfer which involve both spatial and social processes equally. It is therefore important to have a flexible tool to manage the relative importance of these two components of movement patterns, and the multiple criteria analysis of the TOPSIS method provides flexibility in this decision-making process.

Further, integrating the mobility of multiple species into the implementation of protected areas is a major conservation challenge as managers attempt to reach the best compromise between which species to protect and the size of the area which is put under protection. For instance, investigating the movement network of multiple fish species in a multilayer network framework could help managers to identify the most valuable design for delineating a marine protected area that more effectively protects ecosystem functioning. The multilayer network approach might also help measure and mitigate against the spatial and temporal interactions between human and wildlife mobility patterns. A recent review paper (Meekan et al., 2017) highlighted how comparing human and wildlife spatial networks can provide valuable information for anticipating conservation actions. As such, a multilayer network framework will potentially facilitate our predictions of the timing and location of human-wildlife conflict or collisions. Human networks can be inferred from mobile phone data and transport routes of car or ships from global positioning systems while wildlife mobility networks can be determined from various automated monitoring techniques (e.g. camera traps, biologging). Predicting interactions from poacher and animal mobility networks based on multilayer movement networks might further help rangers improve the efficiency of their own movements for tracking down poachers across large areas while predicting probabilities of interaction between vehicles and wildlife might help prevent and mitigate collisions.

Another promising application of multilayer movement networks includes parasite and disease transmission and information diffusion processes. Preliminary investigations have been pushing towards the use of multilayer contact networks to study diffusion processes in populations (including pathogens, disease, parasites or information) (Silk, Drewe, et al., 2018; Finn, Silk, Porter, & Pinter-Wollman, 2019). We aim to contribute to this discussion by proposing a holistic approach linking spatial and social networks, to estimate the probabilities of transmission and/or diffusion within the population. This approach can not only identify individuals that serve as vectors, but also identify the spatial hubs of infection or information spreading. Simulated multilayer node removal experiments might also help to implement efficient strategies in mitigating or managing infection outbreaks. Finally, probabilities of contact within multilayer movement networks might also inform our understanding of predator-prey dynamics, helping to predict the spatial scales of predator-prey interactions and predation risk (Gaynor, Brown, Middleton, Power, & Brashares, 2019).

### Future methodological developments and challenges

While we illustrate the potential of our framework by analysing the centrality of nodes within a multilayer context, there are a broad range of multilayer network metrics already developed, that might be utilised to tackle important ecological questions. For example, centrality variability between layers has been examined using PageRank centralities and versatilities (De Domenico, Solé-Ribalta, Omodei, Gómez, & Arenas, 2015; Finn et al., 2019). This might feasibly be applied to multilayer movement networks to quantify the repetitive importance of a node across layers. Other approaches such as maximum modularity or community detection procedures can also be applied, which would reveal the propensity of a node to belong to distinct communities across layers depending on the presence and weights of the interlayer and intralayer edges. This approach has been already tested in ecological networks to reveal that the relationship between parasitism and herbivory depends strongly on the extent to which processes in one layer affect those in other layers (Pilosof, Porter, Pascual, & Kéfi, 2017). Another interesting avenue for further research would be node removal simulations to explore both the horizontal and vertical robustness of the network to various habitat perturbations, an approach used for monolayer networks (Mourier, Brown, & Planes, 2017).

A challenge we foresee is one that befalls any network data; the violation of the assumption of data independence. In monoplex networks, the use of randomization procedures provides a means to conduct null hypothesis significance testing on the data for robust interpretation of the patterns extracted (Croft, Madden, Franks, & James, 2011; Farine, 2017). However, because they are made of intra and interlayer edges, null models are non-trivial in multilayer networks. For multilayer movement networks made of different individual layers, new techniques for the development of null models will be needed. Movements networks are built from a time series of movements from locations to locations (Jacoby & Freeman, 2016), so steps can be randomized following a data stream permutation procedure for all individual tracks to construct a null model (Spiegel, Leu, Sih, & Bull, 2016; Farine, 2017). However, this approach is likely to be time consuming as it requires rebuilding a multilayer network after each permutation. Another possibility is to conduct permutations of the movement matrices (i.e. intralayer permutation procedure) or alternatively of the social matrices (i.e. interlayer permutation procedure); comparing both null models may help assess the contribution of the social versus the spatial component to the non-random structure of the multilayer movement network. Such an approach has been previously employed in ecological multilayer networks. For example, Timóteo, Correia, Rodríguez-Echeverría, Freitas, & Heleno (2018) tested how the structure of a spatial network is influenced by the seed–dispersal process within each habitat (intralayer null model) and by the identity of the animals connecting these habitats (interlayer null model). Reshuffling interactions within each habitat (intralayer null model) overestimated modularity, whereas reshuffling the identity of animals in each habitat (interlayer null model) underestimated modularity, demonstrating that interlayer strength was more important than intralayer strength. Finally, we call for future work in developing permutation procedures directly on the supra-adjacency matrix which may save time and computational power.

Animal populations or ecosystems are characterised by many interacting individuals that give rise to emergent behaviour and processes. Networks can provide an appealing tool for studying the complexity of these interactions. Real systems are often interconnected, with many interdependencies that are not properly captured by single-layer networks. Animal movements are often underpinned by decisions that are influenced by spatial, social and environmental processes and a multilayer approach is better able to deal with this multifaceted complexity. To account for this source of complexity, we proposed a more general framework, in which movement networks interact with each other.

## ACKNOWLEDGMENTS

We are grateful to Dr. Michelle Heupel for the use of the Blacktip Reef and Grey Reef sharks datasets. The Blacktip Reef and Grey Reef sharks datasets, and Heron Island acoustic receivers information were sourced as part of the Integrated Marine Observing System (IMOS) Animal Tracking facility—IMOS is supported by the Australian Government through the National Collaborative Research Infrastructure Strategy and the Super Science Initiative. The research on Blacktip Reef and Grey Reef sharks movement was conducted under research permits from the Great Barrier Reef Marine Park Authority (G10/33754.1 and G10/33758.1) and its funding was provided as part of a Future Fellowship (#FT100101004) to Michelle R. Heupel from the Australian Research Council; additional funding was provided by the Australian Institute of Marine Science (AIMS). All Blacktip Reef and Grey Reef sharks research was conducted under James Cook University (JCU) Animal Ethics Permit A1566.

## AUTHOR’S CONTRIBUTION

J.M, E.J.I.L and D.M.P.J. conceived the ideas and designed the methodology; E.J.I.L. participated in collecting the data used to illustrate our concept; J.M. and D.M.P.J. analysed the data; All authors contributed critically to the drafts and gave final approval for publication.

## DATA AVAILABILITY

All data and R codes to conduct the main analyses and generate the figures will be publicly available and archived on Dryad Digital Repository (to be deposited if accepted).

